# Distinctiveness and continuity in transcriptome and connectivity in the anterior-posterior axis of the paraventricular nucleus of thalamus

**DOI:** 10.1101/2022.02.13.480207

**Authors:** Yasuyuki Shima, Henrik Skibbe, Yohei Sasagawa, Noriko Fujimori, Itoshi Nikaido, Nobutaka Hattori, Tadafumi Kato

## Abstract

The paraventricular nucleus of the thalamus (PVT) projects axons to multiple areas and mediates a wide range of behaviors. Heterogeneity of functions and axonal projections in PVT have been reported, but what cell types exist in PVT and how different they are have not been addressed. We applied single-cell RNA sequencing to depict transcriptomic characteristics of mouse PVT neurons. The transcriptome of PVT neurons had a continuous distribution with the largest variance corresponding to the anterior-posterior axis. Although the single-cell transcriptome classified PVT neurons into four types, transcriptomic and histological analyses showed their continuity. Similarly, anterior and posterior subpopulations had nearly non-overlapping axon projection patterns, while another population showed intermediate patterns. In addition, they responded differently to appetite-related neuropeptides, and their chemogenetic activation showed opposing effects in food consumption. Our studies showed contrasts and continuity of PVT neurons underlying their function as a behavior-modulating hub.

## Introduction

Accumulating evidence reveals many facets of the paraventricular nucleus of the thalamus (PVT) in behavioral regulations. PVT receives strong peptidergic and monoaminergic inputs from hypothalamus and brainstem and projects to various areas including the nucleus of accumbens (ACB, we follow the nomenclature of the Allen Mouse Brain Common Coordinate Framework version 3: CCfv3, Wang et al., 2020), amygdaloid complex, the bed nucleus of stria terminalis (BST), and medial prefrontal cortex (mPFC) (Kirouac, 2015; Parsons et al., 2007; Sugiyama et al., 2019; Vertes and Hoover, 2008). PVT is involved in multiple behaviors such as fear memory (Do-Monte et al., 2015; Penzo et al., 2015), reward-seeking (Do-Monte et al., 2017; Keyes et al., 2020; Otis et al., 2017; Otis et al., 2019; Zhu et al., 2016), food intake (Horio and Liberles, 2021; Meffre et al., 2019; Zhang and van den Pol, 2017), and sociability (Yamamuro et al., 2020). PVT also mediates emotion (Kasahara et al., 2016; Kato et al., 2019), saliency (Zhu et al., 2018), and wakefulness (Ren et al., 2018) and therefore assumed as a crucial brain network node/hub for motivated behaviors (Iglesias and Flagel, 2021), anxiety (Kirouac, 2021), and behavior homeostasis (Penzo and Gao, 2021).

Several lines of evidence suggest that there are regional differences in functions and axonal projection patterns in PVT, especially between anterior PVT (aPVT) and posterior PVT (pPVT) (Do-Monte et al., 2017; Gao et al., 2020; McGinty and Otis, 2020). Drd2 was shown to be a marker of the pPVT population (Clark et al., 2017). Cre-positive and Cre-negative PVT cells of Drd2-Cre mice have different axonal projection patterns and functions in saliency and wakefulness, and thus the two cell type model has been proposed (Gao et al., 2020). In addition, functionally distinct populations have been identified by the differences in axon projection targets (Ma et al., 2021; Penzo et al., 2015). On the other hand, detailed retrograde tracing studies (Dong et al., 2017 ; Li et al., 2021) revealed extensive heterogeneity of projection targets among neighboring PVT neurons and found no localized subpopulation in PVT (Li et al., 2021).

To obtain comprehensive knowledge of the cellular diversity of PVT, we attempted to classify PVT cells with their gene expression patterns. We performed single-cell RNA sequence (scRNAseq) from PVT tissues and identified PVT neuronal population from their marker gene expressions. PVT neurons were classified into 4 cell types with continuity, and the most striking transcriptomic variance existed along the anterior-posterior axis. With Cre lines labeling limited populations of PVT, we found nearly non-overlapping projection patterns among PVT subpopulations. Finally, we found that the subpopulation of aPVT and pPVT showed different responses to appetite-related peptide hormones and opposingly modulated food intake behavior. Our research revealed the transcriptomic and anatomical basis for the functional heterogeneity of PVT neurons.

## Results

### Classification of PVT cell types with Quartz-seq2

We employed Quartz-seq2 (Sasagawa et al., 2018), which was evaluated as the best in gene detectability among existing scRNAseq methods (Mereu et al., 2020). Mice with Calb2-Cre and EYFP Cre reporter (Ai3, Madisen et al., 2010) were used to delineate the PVT region (Kirouac, 2015, Supplementary Figure1A). We classified 2,069 qualified cells into 15 cell groups with the Louvain algorithm and annotated their cell types with marker gene expressions (Figure 1A and Supplementary Figure1B). Most neuronal cells (all excitatory) were in single clusters in both principal component analysis (PCA) and Uniform Manifold Approximation and Projection (UMAP) dimension reduction plots (Figure 1A). Neurons also had higher numbers of unique molecular identifiers (UMI) and detected genes than non-neuronal cell types (Figure 1C). After selecting neurons to analyze further, we found that some neurons formed clusters with high ratios of transcripts from mitochondria-encoded genes (mt-genes), which is characteristic of “unhealthy” cells (Sasagawa et al., 2018). Indeed, there was a negative correlation between mt-gene rates and the numbers of genes detected (Supplementary Figure1D, Pearson r = -0.491, p = 2.19 × 10^−62^). We also found that the marker genes of some clusters were expressed not in PVT but neighboring regions (Supplementary Figure1E and F, group #4 and 5). After eliminating the two non-PVT cell types and high mt-gene (> 10 %) population, we obtained 605 cells and hereafter call the remaining population core PVT neurons (Figure 1A).

**Figure 1.**
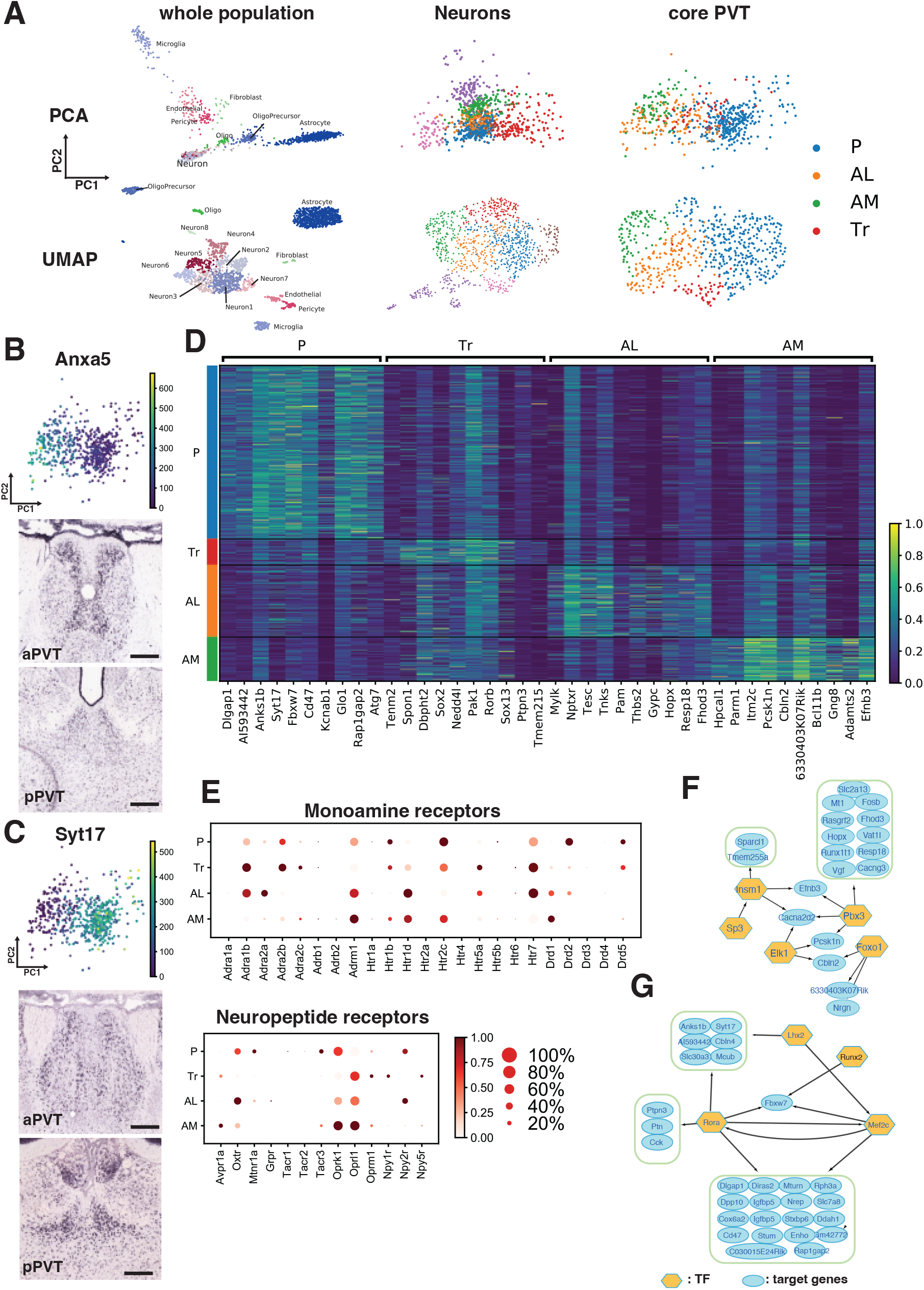
Classification of PVT cells and identification of PVT core cell types. (A) Scatter plots of PCA (top row) and UMAP (bottom row) of the whole population (left), neurons (center), and core PVT cells (right). (B and C) Examples of expression of genes with negative PC1 coefficient (B, Anxa5) and positive coefficient (C, Syt17). Dashed lines: putative PVT area. Scale bars: 200 μm. *in situ* images from mouse.brain-map.org. (D) Heatmap of cell type marker-candidate genes. (E) Dot plots of monoamine and neuropeptide receptors. (F and G) Predicted gene regulatory networks of top DEGs in the anterior (F) and the posterior (G) groups. Genes in rounded green rectangles were regulated by the same TF(s).

### Transcriptomic diversity and continuity in core PVT cells

To ask if there is any practical implication in PCA, we looked into the expression patterns of genes contributing PCs in the Allen Mouse Gene Expression Atlas database. Genes contributing to PC2 (Rheb, Cox7b, Gria2, and Nav1, for example) showed broad expression in PVT (data not shown). However, those with negative PC1 coefficients had strong expression in aPVT while much fewer cells expressed them in pPVT (Figure 1B. Also see Supplementary Figure1G for more examples). In contrast, genes with positive PC1 coefficients, Syt17 for an example, had clear expression patterns in pPVT but had weaker expression in aPVT (Figure 1C and Supplementary Figure1H). Therefore, the axis of the largest variance in gene expression appeared to have topographical correspondence.

Next, we sought an appropriate condition for classifying the core PVT cells. The Louvain algorithm with the default parameter generated nine cell groups (Supplementary Figure1I). However, most cell type marker candidates were expressed in multiple cell types, and few cell types could be distinguished from others by their gene expression (Supplementary Figure1J). Therefore, we applied a parameter of the Louvain algorithm in which all cell types had unique marker genes and then obtained four cell types (Figure 1A and D). The most abundant population’s marker genes were strongly expressed in pPVT and thus named posterior (P). The second and the third groups were named anterior lateral (AL) and anterior medial (AM) from their marker gene expression (see Figure. 2). The smallest population expressed both anterior and posterior genes and was thus named transitional (Tr) cells.

**Figure 2.**
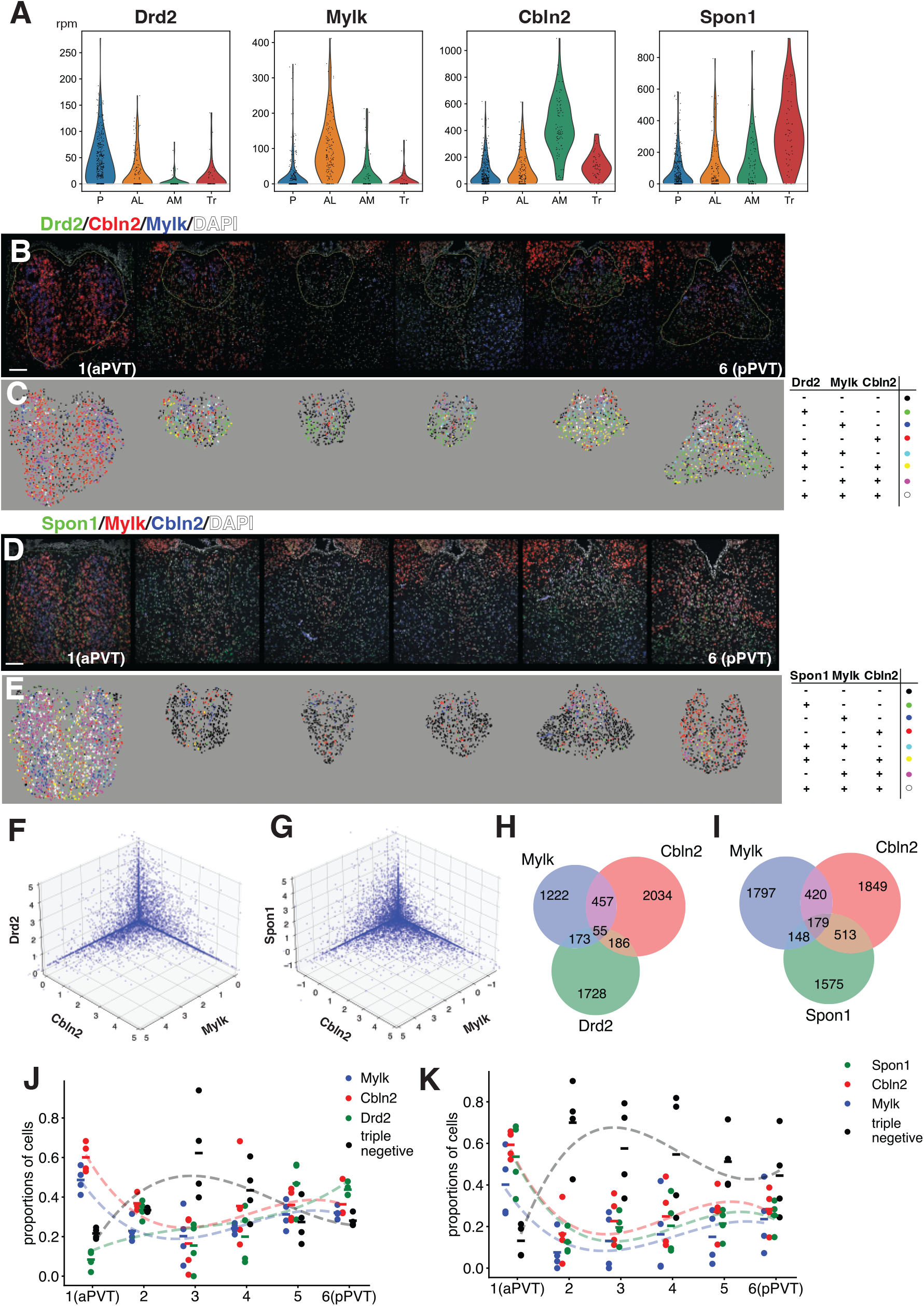
*In situ* hybridization of cell-type marker genes. (A) Violin plots of markers of each cell type. (B) An example of a set of 6 sections from anterior (left) to posterior (right) PVT stained with Drd2 (green), Cbln2 (red), Mylk (Blue), and DAPI (white). Yellow lines: areas used for signal quantification. (C) Marker positive cells were colored with the expression code (right). (D-E) Triple fluorescent *in situ* images (D) of Spon1 (green), Cbln2 (red), Mylk (blue), and DAPI (white) and distribution of marker-positive cells. (F and G) Scatter plots of clipped signals in Drd2/Cbln2/Mylk (F) and Spon1/Cbln2/Mylk *in situ* (G). (H and I) Venn diagrams for strongly marker positive (++) cells in Drd2/Cbln2/Mylk (H) and Spon1/Cbln2/Mylk (I)*in situ* (J and K) proportions of marker-positive (+) cells in each anterior-posterior level for Drd2/Cbln2/Mylk (J) and Spon1/Cbln2/Mylk (K) *in situ* hybridization. Lines: averages of each data point. Dotted lines: cubic spline interpolation. Scale bars: 100 μm.

The four cell types were not discrete but continuous populations. A partition-based graph abstraction (PAGA, Supplementary Figure1K) showed the connections between all pairs of cell types. As expected, visualization of continuity of cell groups with diffusion pseudotime rank analysis (Stanley et al., 2020) displayed the existence of intermediate population between any pairs of cell types (Supplementary Figure1L). These results demonstrated both prominent differences and continuity between aPVT and pPVT.

### Differentially expressed genes and transcriptional networks

PVT receives extensive and various neuromodulatory inputs, and regional differences of their effects have been suggested (Kirouac, 2015, 2021; McGinty and Otis, 2020). Indeed, we found the four cell types have characteristic expressions of their receptors (Figure 1E. See also Figure 4A). Next, we investigated what types of genes are differentially expressed between anterior (AL and AM) and posterior (P) groups. We identified 366 differentially expressed genes (DEG, FDR < 0.01, log fold changes > 0.25, Supplementary Table1). Enrichment analysis of Gene Ontology terms of DEG indicated expression differences in several neuronal functions, such as synaptic signaling, regulation of ion transport, and axon development (Supplementary Table2). We analyzed gene regulatory networks of the DEGs with pySCENIC (Van de Sande et al., 2020). Pbx3 regulated the largest number of anterior DEGs, and some DEGs were regulated by multiple key transcriptional factors (TFs) in the anterior group (Figure 1F). The posterior network indicated mutual regulations between Rora and Mef2c, and the TF pair regulated most of the posterior DEGs (Figure 1G).

### Cell type marker distributions in PVT

To examine the distribution of each cell type, we looked for “marker” genes that were selectively expressed in one of the cell types and able to distinguish from others by *in situ* hybridization. While many marker gene candidates had substantial expression in other cell types, we chose Drd2, Mylk, Cbln2, and Spon1 as marker genes for P, AL, AM, and Tr, respectively (Figure 2A). We performed multicolor fluorescent *in situ* hybridization with RNAscope multifluorescent kit (Supplementary Figure2A). The markers of aPVT showed strong expression in most anterior sections, while posterior sections had a broad expression of P marker Drd2 (Figure 2B, C). In anterior-most PVT sections, Cbln2 positive cells formed columnar clusters along the midline, while Mylk positive cells laterally aligned to the Cbln2 clusters. As reported previously (Gao et al., 2020) anterior markers (Mylk and Cbln2) were also expressed in dorsal pPVT. All AL and AM marker candidates had expression in dorsal pPVT (for example, see Supplementary Figure1G), and our transcriptome analysis could not distinguish AM/AL population in pPVT and those in aPVT.

We set thresholds for marker-positive (+) cells and strongly positive (++) cells from the distribution of Z-score normalized signals (Supplementary Figure2B). Marker-positive cells were painted with the color code shown in Figure 2C (also see marker ++ cells’ distribution in Supplementary Figure2C). The expression of Tr marker Spon1 was also visualized in the same way (Figure 2D-E and S2E). Spon1 positive cells were mainly found around the midline of anterior-most PVT sections. The scatter plots of threshold-clipped signals (Figure 2F and G, see S2C for a plot before clipping) and Venn diagrams of strongly positive cells (Figure 2H and I) demonstrated the separation of PVT population by the expression of the marker genes.

Next, we investigated the distribution of marker-positive cells along the anterior-posterior axis (Figure 2J-K). As expected, the rates of Cbln2- or Mylk-positive cells were highest in aPVT, and posterior cell type marker (Drd2) was highly expressed in pPVT. On the other hand, sections in the middle (#3 and #4) had high proportions of cells with no marker gene expression (triple negative, black in Figure 2J). The Tr marker Spon1-positive cells were enriched in the anterior-most sections, and the middle sections had high proportions of triple-negative cells also in the Spon1/Cbln2/Mylk *in situ* (Figure 2K). Another set of cell type markers (Gng8, Ecel1, and Col12a1 for AM, AL, and P, respectively) also showed similar tendency (Supplementary Figure2F-H). Thus, the substantial number of neurons in central PVT do not have strong expression of any of the cell type markers. By contrast, we found that genes expressed in all core PVT cell types showed consistent A-P distributions. Efnb3, Maoa, and Maob have restricted expression in PVT but expressed in all cell types (Supplementary Figure2I–J), and the proportions of gene expression-negative cells were low in middle sections (Supplementary Figure2K). We also investigated the expression of marker genes for the finer cell type classification (Supplementary Figure1J) in the Allen *in situ* database to examine if specific cell type(s) for central PVT existed, but all of the candidate marker genes had strong expression either in aPVT or pPVT (data not shown). These results indicated that transcriptionally distinctive populations are mainly distributed in aPVT and pPVT, while the majority of central PVT neurons have less characteristic, intermediate gene expression patterns.

### Distinct axonal projections of genetically defined subpopulations

To compare axonal projections of specific cell groups, we searched for mouse Cre lines with restricted expressions in PVT cell types. Among Cre lines characterized by the Allen Mouse Connectivity project (Oh et al., 2014), we selected Ntrk1-Cre and Drd2-Cre as candidates. Ntrk1 is expressed in both AL and AM populations (Supplementary Figure3A), and multifluorescent *in situ* showed that Ntrk1 was mostly co-expressed with either/both Cbln2 and Mylk (Supplementary Figure3B and C). The number of Ntrk1-positive cells was smaller than those of Cbln2 and Mylk (Supplementary Figure3D), thus Ntrk1-positive cells are mostly subpopulations of AL and AM. We also examined Cre expression in Ntkr1-Cre mice and found similar co-expression patterns (Supplementary Figure3E and F).

To characterize axonal projection patterns of these two lines, we injected AAV encoding EGFP Cre reporter (AAV-CAG-FLEX-EGFP) at aPVT and pPVT of both lines. Since none out of 10 Drd2-Cre animals injected at aPVT had reporter expression, we concluded that no cells in aPVT of Drd2-Cre expressed Cre strongly enough to drive the AAV reporter. Two brains from each condition were imaged with TissueCyte serial sectioning tomography to reconstruct axonal projections in the whole brain. Taken images were mapped to CCFv3 (Wang et al., 2020) to make whole-brain 3-D models (Supplementary Figure3G and Supplementary Video 1).

Ntrk1-Cre labeled aPVT cells (aPVT^Ntrk1^) and Drd2-Cre labeled pPVT cells (pPVT^Drd2^) showed distinctive, almost non-overlapping axonal projection patterns (Figure 3A). In the frontal cortex, pPVT^Drd2^ projects to the prelimbic area (PL) and the agranular insular area, dorsal part (AId), while axons from aPVT^Ntrk1^ were found in the infralimbic area (ILA). In the striatum (STR), aPVT^Ntrk1^ projection was restricted to the mediodorsal ACB shell, while neurons in pPVT^Drd2^ projected widely to the ventral part of STR, including the olfactory tubercle (OT). In the amygdaloid body, both Ntrk1-aPVT and Drd2-pPVT projected to the central amygdala nucleus (CEA), and aPVT^Ntrk1^ had strong axonal projections in the basomedial amygdala nucleus (BMA). BST received strong input from aPVT^Ntrk1^ while there are few axons from pPVT^Drd2^. Instead, projections from pPVT^Drd2^ were observed in the fundus of the striatum (FS). Dominant projections from aPVT^Ntrk1^ were also seen in the hypothalamus (HY) and subiculum (SUB, arrows in Supplementary Figure2G). The whole-brain level projection images demonstrated projection pattern differences between aPVT^Ntrk1^ and pPVT^Drd2^ (Figure 3B). We quantified axonal signals on each brain structure in the 3D model and calculated each structure’s proportion of the signals (see Methods). Figure 3C displays proportions of axon projections in brain structures with more than 1 % of axonal signals. As shown in the left column, most structures had more than twice (|log_2_ ratio| > 1) differences in proportions (represented by red or blue numbers in Figure 3C), supporting apparent differences of axonal projection patterns between aPVT^Ntrk1^and pPVT^Drd2^.

**Figure 3.**
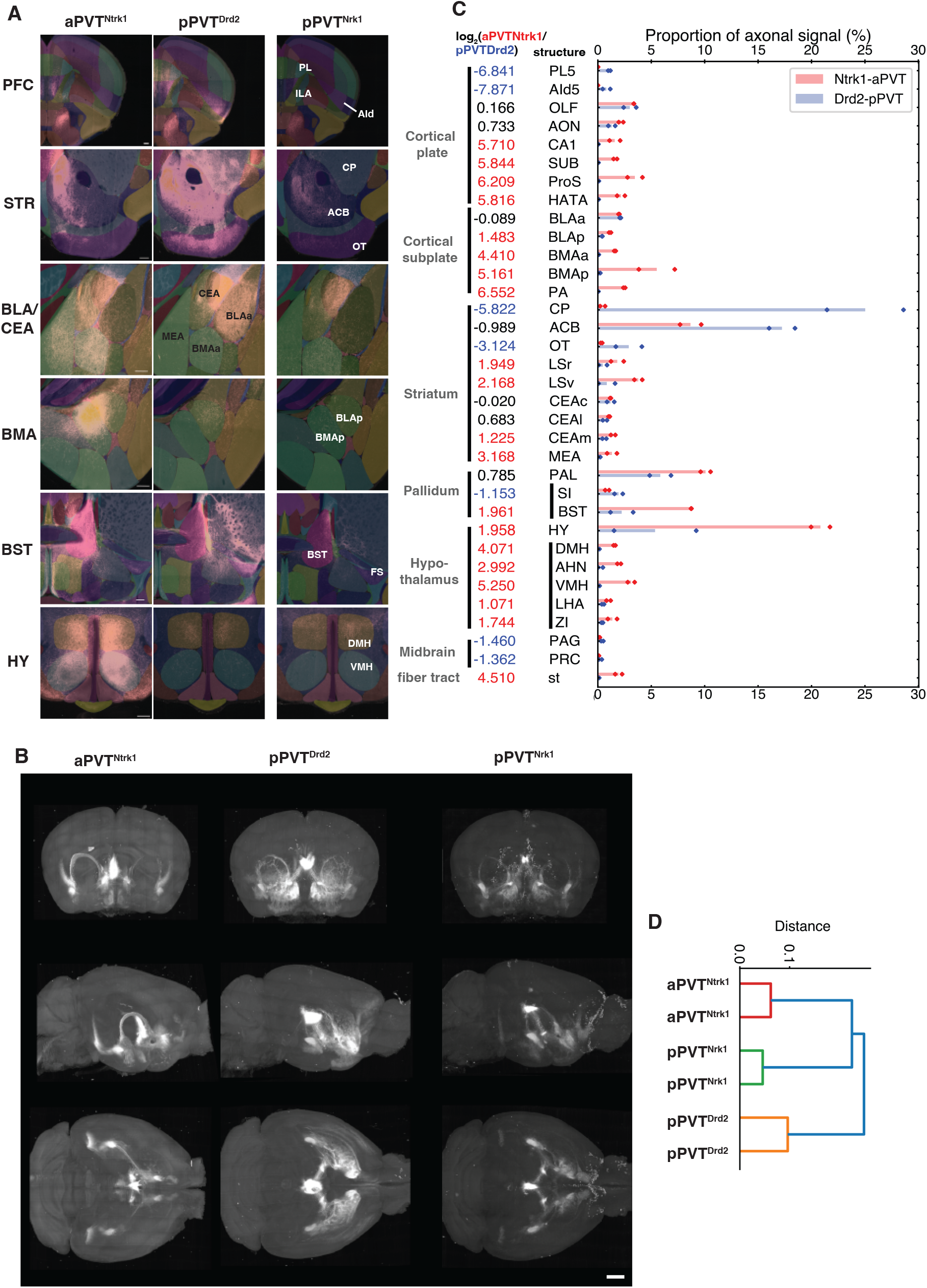
Whole-brain reconstruction of subpopulation-specific axonal projection. (A) Representative brain regions with differential axonal projections. (B) Coronal (top row), sagittal (middle row), and horizontal (bottom row) projections of images from aPVT^Ntrk1^(left column), pPVT^Drd2^ (center column), and pPVT^Ntrk1^ (right column). (C) The proportion of axonal signals in each brain region with more than 1 % of projections. Log ratios of aPVT^Ntrk1^ and pPVT^Drd2^ are shown on the left. Colors of number represent dominance (|log_2_ ratio| >1) of axonal projections of aPVT^Ntrk1^ (red) or pPVT^Drd2^ (blue). (D) A dendrogram of axonal projection patterns. See Supplementary Figure3 for abbreviations of brain structures. Scale bars: 100 μm (A) and 500 μm (B).

Although the number of infected neurons was much smaller, pPVT^Ntrk1^showed consistent reporter expression, and its axonal projection patterns share the characteristics with those of both aPVT^Ntrk1^ and pPVT^Drd2^. Striatal projections were seen in both the shell and core of ACB. Amygdala projection was mainly observed in CEA and few in BMA. In BST, only the lateral part received the projection like pPVT^Drd2^, while they have SUB projection like aPVT^Ntrk1^(arrows in Supplementary Figure4H). A dendrogram of projection data supported that pPVT^Ntrk1^had a distinct projection pattern from aPVT^Ntrk1^ and pPVT^Drd2^ (Figure 3D).

### Differential responses to neuropeptides

PVT receives inputs from hypothalamic appetite-related neuropeptides (Kirouac, 2015). Orexin receptor Hcrtr1 and Hcrtr2 were expressed in anterior cell types while melanocortin-stimulating hormone (MSH) receptor Mc3r’s expression was dominant in P cell type (Figure 4A). To investigate responses of PVT neurons to these two appetite-related hormones, we infected the GFP Cre reporter AAV either at aPVT of Ntrk1-Cre or pPVT of Drd2-Cre and performed whole-cell patch-clamp recording from GFP-labeled neurons in acute slices of the brains. Most thalamic neurons including PVT (Zhang et al., 2009) spike after hyperpolarization (low threshold spike: LTS), and both aPVT^Ntrk1^and pPVT^Drd2^spiked once or twice after hyperpolarization (Figure 4B and C). After applying Orexin-A (OrxA), we found the number of LTS increased significantly in aPVT^Ntrk1^. pPVT^Drd2^ did not show such an increase of LTS with OrxA application but had a similar elevation of LTS with γ-MSH, a ligand for Mc3r (Figure 4B and C. See Supplementary Figure4A for input-frequency curves). aPVT^Ntrk1^and pPVT^Drd2^showed similar ligand-specific responses in the depolarization of resting membrane potentials (Figure 4B and D) and firing frequency by current injection (Supplementary Figure4A).

**Figure 4.**
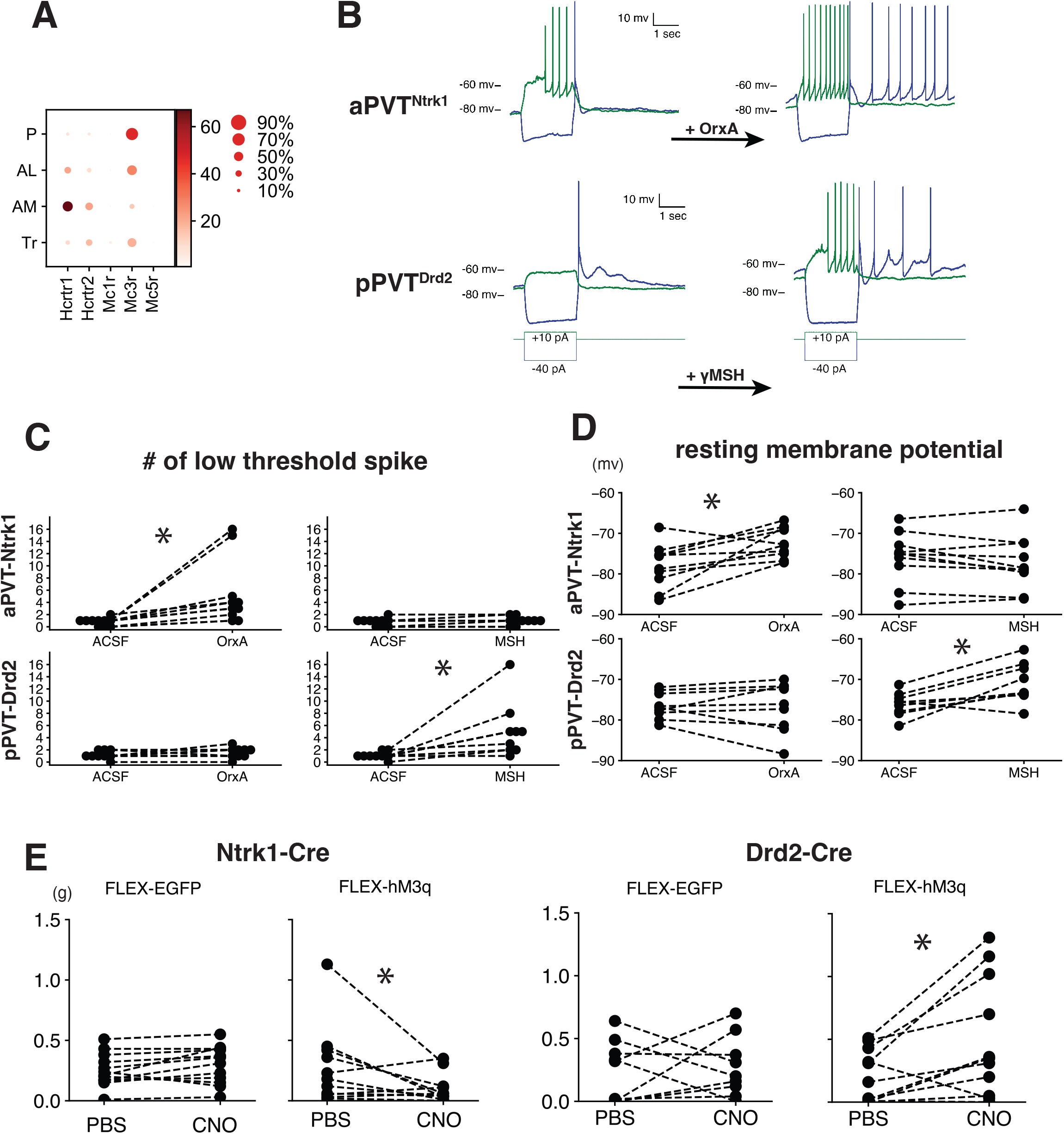
Physiological and behavioral differences between aPVT^Ntrk1^ and pPVT^Drd2^. (A) Dot plots of orexin and melanocortin-stimulating hormone receptors. Mc2r and Mc4r were not detected. (B) Firing patterns of PVT neurons upon current injection. Top: traces from aPVT^Ntrk1^ neuron before (left) and after (right) or an application. Bottom: traces from a pPVT^Drd2^ neuron before (left) and after (right) γMSH application. (C) Numbers of low-thresholds spikes in 3 seconds after hyperpolarization. (D) Resting membrane potential before and after ligand application. (E) The amount of consumed food in 3 hours after PBS or CNO injection. Asterisks: p < 0.05 with paired Wilcoxon rank test after Bonferroni correction. n=10 in each condition in C and D. In F, four males and four females were used in each condition.

### Differential effects in food consumption by chemogentic activation

Physiological recordings suggested that aPVT^Ntrk1^and pPVT^Drd2^could have different roles in appetite regulation. To investigate cell type-specific roles of PVT neurons in food consumption behaviors, we infected Cre reporter AAV encoding EGFP (AAV-CAG-FLEX-EGFP) or hM3q (AAV-CAG-FLEX-hM3q), an activating designer receptor exclusively activated by designer drug (DREADD), in aPVT and pPVT of Ntrk1-Cre and Drd2-Cre, respectively. Infected mice were housed in individual cages and habituated. Mice had an intraperitoneal injection of PBS or clozapine-N-oxide/PBS, and the consumption of food was measured 3 hours after injection. Flex-EGFP encoding AAV injected mice did not show a significant difference in food consumption, but hM3q-AAV injected Ntrk1-Cre mice showed decreased food consumption with CNO injection (Figure 4E). In contrast, hM3q-AAV injected Drd2-Cre mice ate significantly more chows with CNO (Figure 4E).

We also examined if animals showed any preference to activation of either aPVT ^Ntrk1^ or pPVT ^Drd2^ with real time place preference tests as previously shown (Do-Monte et al., 2017; Zhu et al., 2016). We infected AAV ChR2 Cre reporter to the Cre mice, implanted optical cannulas, and tested if infected mice preferred to rooms with or without laser stimulations. Some mice showed strong preference to either laser-on or -off rooms, but there was no statistically significant difference overall (Supplementary Figure4B and C).

## Discussion

Our scRNAseq analysis revealed diversity and continuity in the transcriptome of core PVT neurons. We found the largest variance of gene expression lied in the AP axis of PVT, which supports the previously suggested differences between aPVT and pPVT. Although the transcriptomic contrast exists between aPVT and pPVT, PVT neurons’ transcriptome have a continuous distribution rather than discrete groups. Thalamic neurons have transcriptomic profiles with continuous variations corresponding with their topographical arrangement (Phillips et al., 2019; Roy et al., 2022). Our findings demonstrated that such a topography –gene expression relationship also exists within PVT.

We revealed the existence of subpopulations with nearly non-overlapping axonal projection patterns by genetically selecting highly distinct PVT subsets. Similarly, using specific Cre lines for AL, AM, and Tr (Spon1-Cre (Zhang et al., 2019), for example) would allow more detailed dissection of PVT circuits. However, one of the limitations of this strategy is that it is applicable only to populations with unique marker expression. Thus, it is challenging to specifically label particular intermediates, which can be the origin of extensive divergence of projections of PVT neurons (Li et al., 2021). Also, we showed pPVT^Ntrk1^ had different projection patterns from aPVT^Ntrk1^ and pPVT^Drd2^. This result indicates intermediates also exist in axonal projection patterns. In addition, posterior subpopulations of AL/AM (labeled by pPVT^Ntrk1^) and anterior AL/AM (aPVT^Ntrk1^) were indistinguishable in our transcriptome analysis. Therefore, PVT neurons’ projection patterns (and possibly functions) might depend on both their transcriptomic cell types and spatial locations. For example, orexin injection in pPVT promotes hunger (Meffre et al., 2019) while we showed activation of orexin-receptor expressing aPVT^Ntrk1^ suppress food intake. This apparent contradiction might be explained by the projection difference between aPVT^Ntrk1^ and pPVT^Ntrk1^.

PVT receives various monoamine and neuropeptide inputs from the brainstem and hypothalamus, and our gene expression analysis revealed that the core PVT cell types differentially expressed those neurotransmitter receptors. Indeed, aPVT^Ntrk1^ and pPVT^Drd2^ were activated by different appetite-related peptide hormones, and their chemogenetic activation had opposite effects on food consumption. Together with axonal projection differences, such neuromodulations might affect behaviors by shifting activities of PVT subpopulations.

## Acknowledgment

We are grateful to Keisuke Fukumoto, Yoshimi Iwayama, and Kenji Ohtawa at Research Resource Division, RIKEN Center of Brain Science for their technical assistance, Takaoki Kasahara for critical reading of the manuscript, and Shigeyoshi Itohara and Yuki Kobayashi for supplying materials. This research was supported by Science Grant-in-aid (KAKENHI, Grant No. 17H01573 to T.K. and 20K06919 to Y.Shima.), Brain & Behavior Research Foundation (NARSAD young investigator grant, Grant No. 28168 to Y.Shima), and the Japan Agency for Medical Research and Development (AMED, Grant No.M6021027 to T.K., I.N., and Y.Shima). This research was also supported by the program for Brain mapping by Integrated Neurotechnologies for Disease Studies (Brain/MINDS) from AMED (JP21dm0207001).

## Author contribution

Conception: Y.Shima and T.K. histology: N.F and Y.Shima, sequencing: Y.Sasagawa, I.N., and Y.Shima, tomography: H.S, N.F., and Y.Shima, physiology and behavior: Y.Shima, supervising the research: H.N. and T.K.. All authors contributed to writing the manuscript.

## Figure legends

**Supplementary Figure1.**
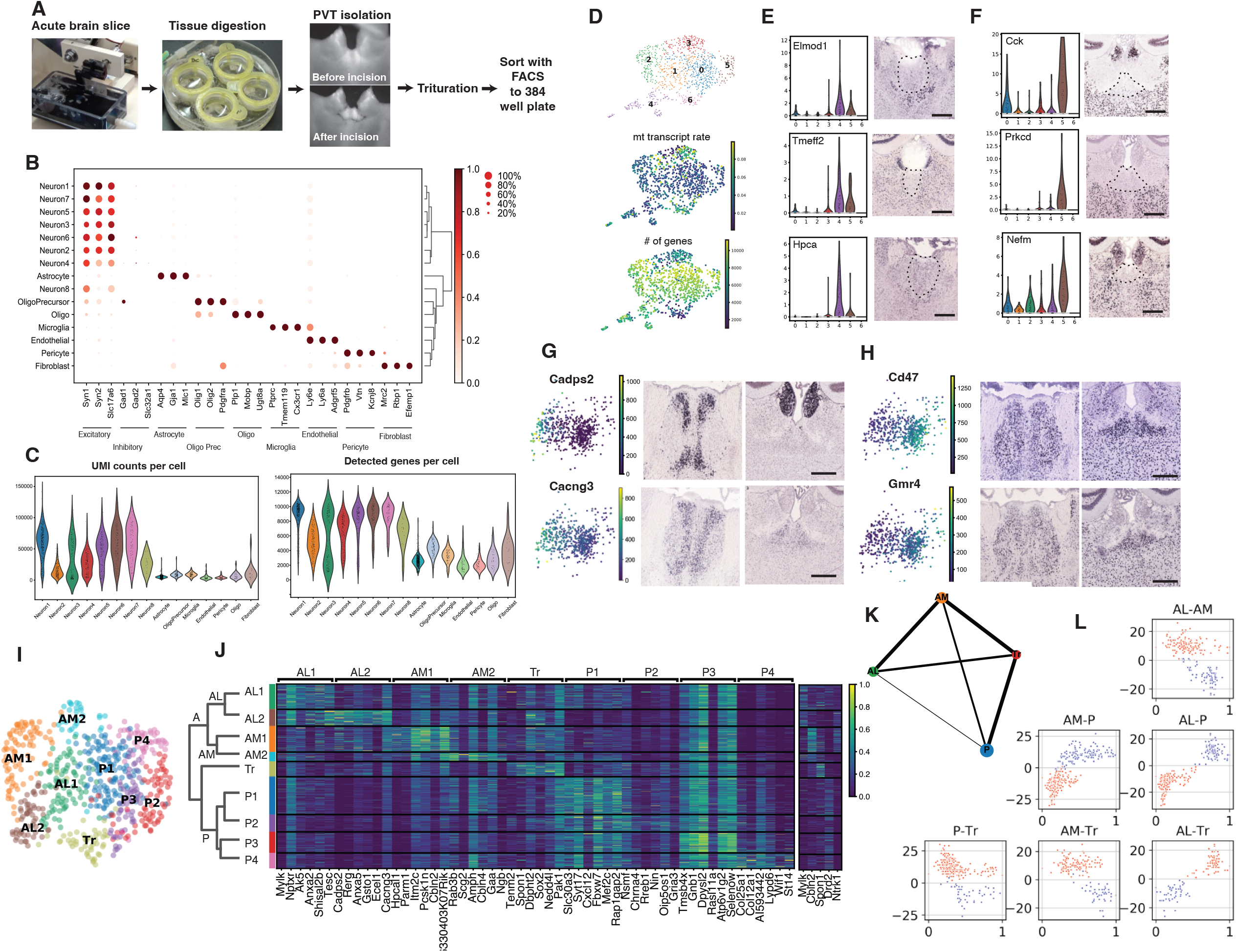
(A) Single-cell isolation method. Brains from Calb2-Cre::Ai3 were sliced, then the slices were incubated with protease-containing ACSF. PVT regions were dissected from slices under the guide of fluorescence. Dissected tissues were triturated with Pasteur pipettes. The dissociated cells were sorted with FACS. (B) Dot plots of major cell type markers in the whole PVT populations. (C) Violin plots of the number of UMI (left) and detected genes (right) per cell. (D) UMAP (top), the rates of mitochondrial transcripts (middle), and the number of genes detected (bottom) of the neurons. (E and F) Marker expression of cell type #4 (E) and #5 (F). dotted area: putative PVT. (G and H) Expression patterns of genes with negative (G) and positive (H) coefficient of PC1. Left: heat map of gene expression on PCA plot, center: *in situ* images of PV, right: *in situ* images of pPVT. *In situ* images in E-H were taken from mouse.brain-map.org (Allen Institute). Scale bars: 300 μm. (I) UMAP of the fine classification of PVT neurons. (J) Heatmap of cell type marker candidates. (K) PAGA graph of 4 cell types. (L) Plots of diffusion pseudotime rank (Stanley et al., 2020) displaying connectivity between each cell type. Diffusion pseudotime (x axis), calculated from the expression level of top 10 differentially expressed genes, is plotted against the relative cluster connectivity (y axis), which was computed by a neighborhood graph.

**Supplementary Figure2.**
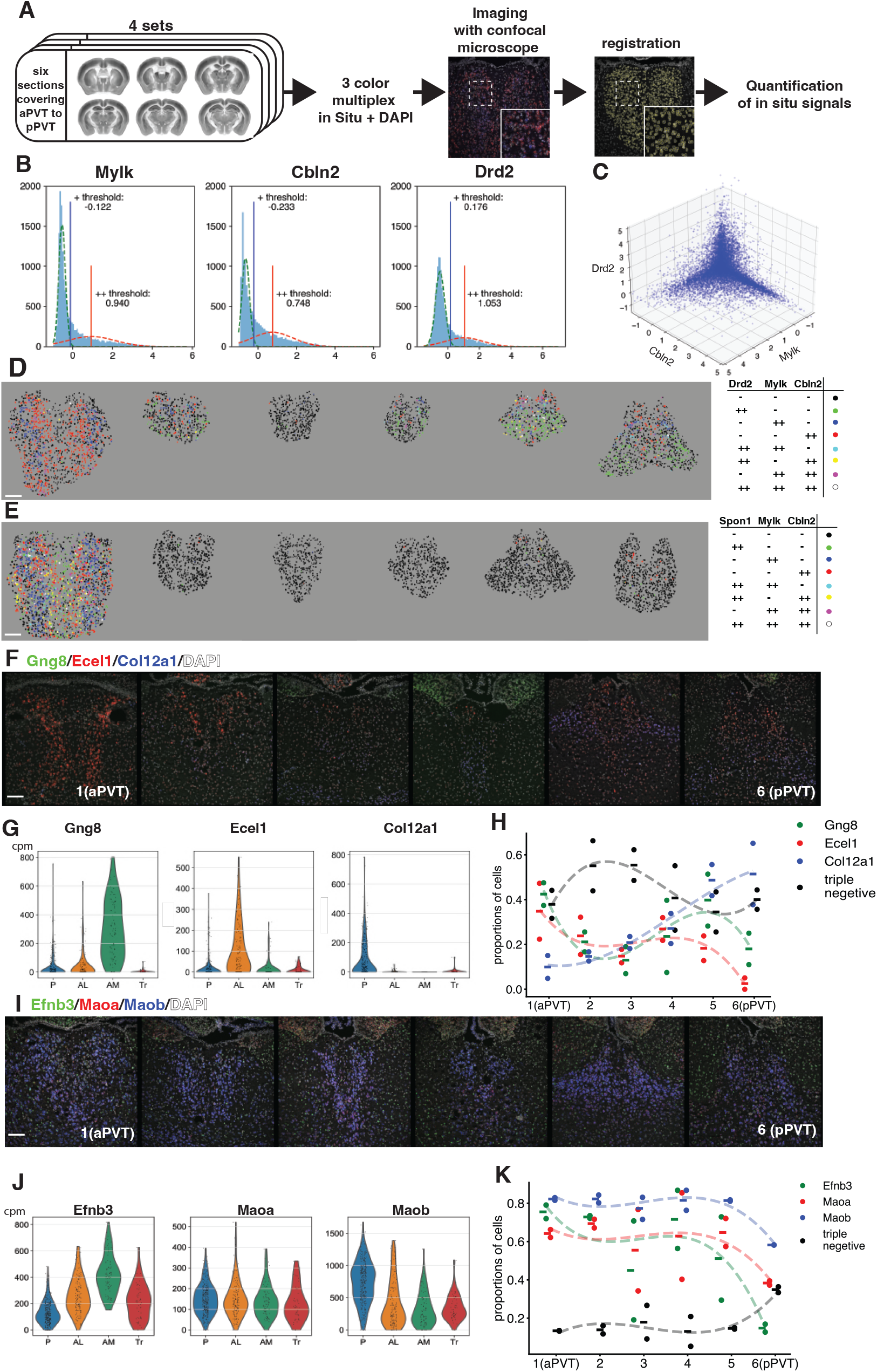
(A) The method of quantification of marker gene expression. We prepared sets of 6 sections covering from aPVT to pPVT on a single slide glass and stained 4 sets with DAPI and the probes of the markers. Images were taken with a confocal microscope, and the mean signal intensities were measured. Insets: magnification of dotted rectangles. Coronal brain section images are from CCFv3. (B) Histograms of *in situ* signal intensities normalized by Z-score. We assumed that each histogram was composed of two Gaussian distributions, a noise component and a signal component, and set a threshold for positive (+) cells where the two curves crossed. Thresholds for strongly positive (++) cells were set at the peaks of signal Gaussian curves. Green dotted lines: Gaussian fit to “noise” component, red dotted lines: Gaussian fit to “signal” component, blue lines: thresholds for marker positive cells (+). red lines: thresholds for strongly marker positive cells (++). (C) A 3D scatter plot of Z-scores of signal intensities of Drd2, Mylk, and Cbln2 before clipping. (D and E) Distribution of ++ cells for Drd2, Mylk, and Cbln2 staining (D) and Spon1, Mylk, Cbln2 staining (E). (F) *In situ* images for Gng8 (green), Ecel1 (red), and Col12a1 (blue) with DAPI (white). (G) Expression of Gng8, Ecel1, Col12a1 in each cell type. (H) A dot plot for proportions of gene-expression positive cells in Gng8, Ecel1, Col12a1 *in situ*. n = 2. (I) *In situ* images for Efnb3 (green), Maoa (red), and Maob (blue) with DAPI (white). (J) Violin plots of genes in core PVT cell types for Efnb3, Maoa, and Maob. (K) Histograms of Z-score normalized fluorescent *in situ* signals for Efnb3, Maoa, and Maob. (L) A dot plot for proportions of gene-expression positive cells in Efnb3, Maoa, and Maob staining. Scale bars: 100 μm.

**Supplementary Figure3.**
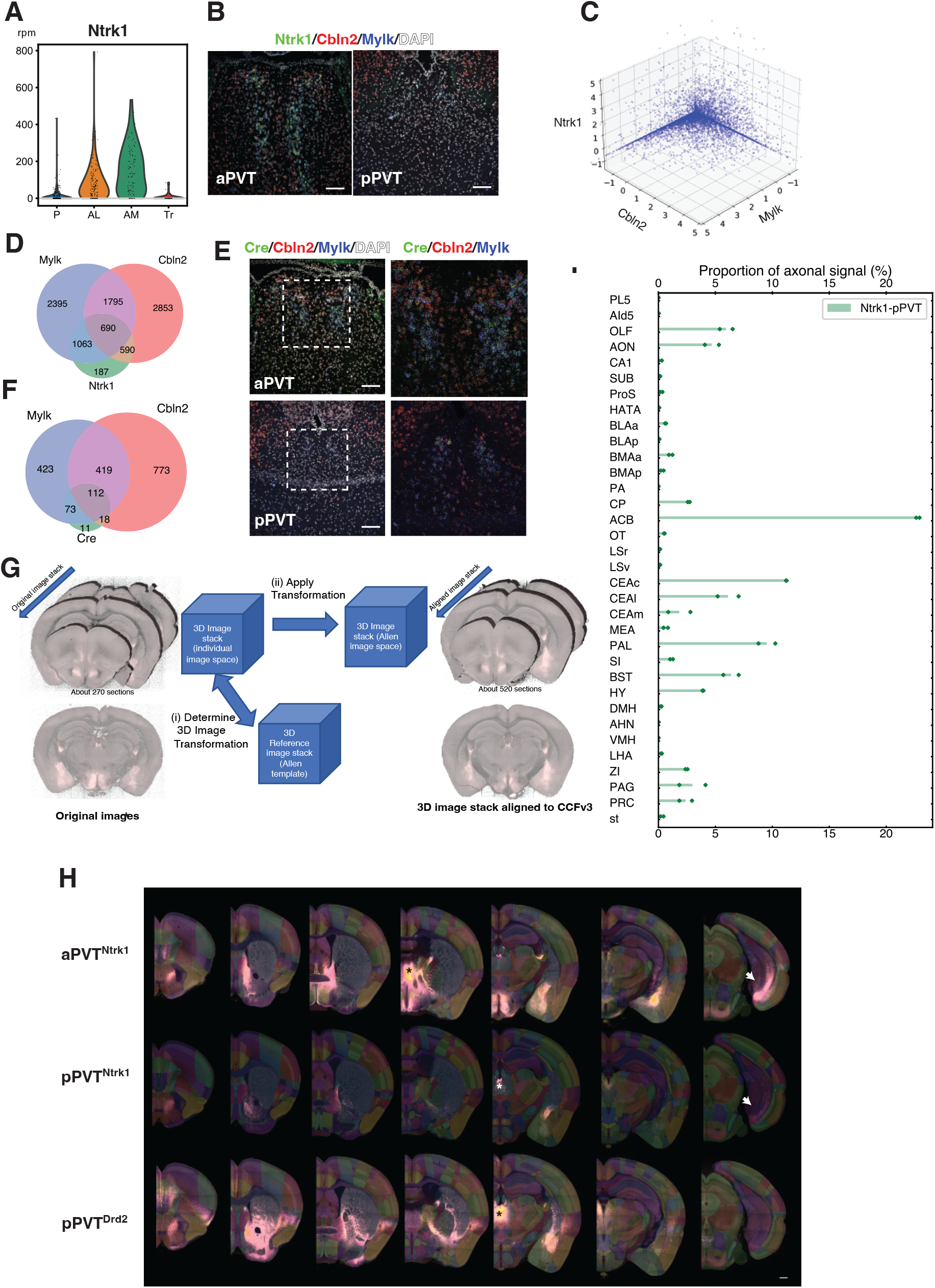
(A) A violin plot of Ntrk1 expression in core PVT cell types. (B) Multicolor in situ of Ntrk1 (green), Cbln2 (red), and Mylk (blue). Right: aPVT, left: pPVT. Scale bars: 100 μm. (C) A plot of signal intensities. *In situ* signals were transformed to Z-score and clipped to thresholds. (D) A Venn diagram of overlaps of Ntrk1-, Mylk-, and Cbln2-positive cells. (E) Multicolor *in situ* of Cre (green), Cbln2 (red), and Mylk (blue). Right panels are magnified images of dotted rectangles in the left panels. Scale bars: 100 μm.(F) A Venn diagram of Cre-, Cbln2-, and Mylk-positive cells. (G) Mapping images to CCFv3. (I) Proportion of axons from pPVT^Ntk1^. The first step determines the spatial transformation that maps the original image stack to the Allen Mouse Brain Common Coordinate Framework based on channel 1. (ii) The second step maps all remaining color channels to the Common Coordinate Framework. (H) Coronal images of Cre-reporter AAV injected brains. Asterisks: virus injection sites, arrows: SUB, scale bar= 300 μm. (I) Proportions of axons from pPVT^Ntrk1^. Abbreviations not shown in the main text are followings: AHN: anterior hypothalamic nucleus, AId5: AI, layer 5, BLA: basolateral amygdala nucleus, BLAa: BLA, anterior part, BLAp: BLA, posterior part, BMAa: BMA, anterior part, BMAp: BMA, posterior part, CA1: Field CA1, CEAc: CEA, capsular part, CEAl: CEA, lateral part, CEAm: CEA, medial part, CP: caudate putamen, DMH: dorsomedial nucleus of hypothalamus, HATA: hippocampal-amygdala transition area, LHA: lateral hypothalamic area, LSr: lateral septal nucleus, rostral part, LSv: lateral septal nucleus, ventral part, MEA: medial amygdala nucleus, mPFC: medial prefrontal cortex, OLF: olfactory areas, PA: Posterior amygdala nucleus, PAG: periaqueductal gray, PAL: pallidum, PL5: Prelimbic area, layer 5, PRC: precommissural nucleus, ProS: Prosubiculum, SI: substantia innominata, st: stria terminalis, VMH: ventromedial hypothalamic nucleus, ZI: zona incerta.

**Supplementary Figure4.**
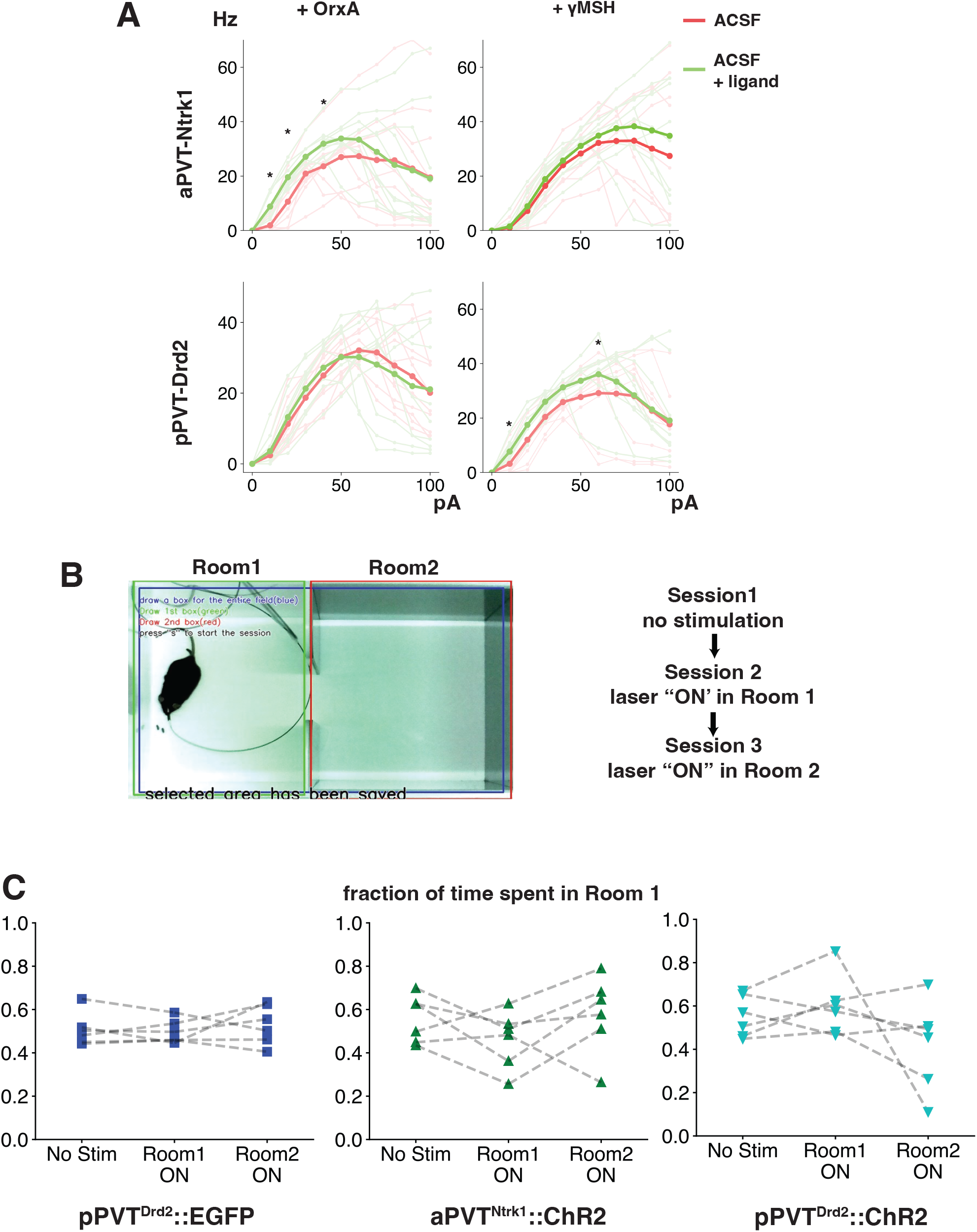
(A) Firing frequency upon current injection. Current clamp recordings were performed with ACSF (red lines), then the same set of recording was performed with bath application of one of the ligands (green lines). Thin lines: Each cell’s average frequencies of sweeps. Thick lines: averages of cells. Asterisks: p < 0.05 with paired sample t-test after Bonferonni correction. n = 10 each. (B) Real-time preference test. Right panel: a screen shot from an experiment. (C) Plots of fractions of time mice spend in Room 1. n = 6 each.

**Supplementary Table 1**

The list of differentially expressed genes between anterior (AL/AM) and posterior (P) PVT.

**Supplementary Table 2**

The list of enriched gene ontology terms.

**Supplementary Video1**

An animation of coronal sections. Maximum signals from all samples were superimposed. Red: aPVT^Ntrk1^, Green: pPVT^Ntrk1^, and Blue: pPVT^Drd2^.

## Method

### Experimental model

Calb2-Cre (B6(Cg)-Calb2tm1(cre)Zjh/J, Taniguchi et al., 2011) and Ai3(B6.Cg-*Gt(ROSA)26Sor*^*tm3(CAG-EYFP)Hze*^/J, Madisen et al., 2010) were obtained from Jackson laboratory. Ntrk1-Cre (B6;129S4-*Ntrk1*^*tm1(cre)Lfr*^/Mmucd) and Drd2-Cre(B6.FVB(Cg)-Tg(Drd2-cre)ER44Gsat/Mmucd, Gong et al., 2007) were obtained from Mutant Mouse Resource & Research Center. All lines were maintained by crossing with wild type C57Bl/6J (Clea Japan, Inc.). All procedures were approved by the Wako Animal Experiments Committee, RIKEN.

### Statistical analysis

edgeR (Chen et al., 2016) was used for statistics in differential gene expression analysis. Otherwise, scipy was used for statistical analysis.

### Brain slice preparation (both scRNAseq and electrophysiology)

We followed slice preparation protocols for aged mice (Tasic et al., 2018; Ting et al., 2018). Mice were deeply anesthetized with isoflurane inhalation and perfused with ice-cold NMDG-ACSF (93 mM N-methyl-D-glucamine, 2.5mM KCl, 1.2mM NaH_2_PO_4_, 20 mM HEPES, 30 mM sodium carbonate, 5 mM sodium ascorbate, 2 mM thiourea, 3 mM sodium pyruvate, 12 mM N-acetylcysteine, 25 mM glucose, pH: 7.3-.4, osmolality: 310 mM). The brain was embedded in 2 % low melting temperature agarose/NMDG ACSF and sliced at 300 µm thick with VF300-0Z compresstome (Precisionary instruments). Cut slices were incubated in warmed (33-34°C) NMDG-ACSF while NaCl spikes (Ting et al., 2018) were applied for physiology experiments. Slices were transferred to HEPES-ACSF (92 mM NaCl, 2.5 mM KCl, 1.2mM NaH_2_PO_4_, 20 mM HEPES, 30 mM sodium carbonate, 2 mM CaCl, 2 mM MgSO4, 5 mM sodium ascorbate, 2 mM thiourea, 3 mM sodium pyruvate, 12 mM N-acetylcysteine, 25 mM glucose, pH: 7.3-.4, osmolality: 310 mM). Both ACSF were bubbled with 95 % O_2_ and 5 % CO_2_ gas. For Quarz-seq2, 1 µM TTX, 50 µM D-AP5, and 20 µM CNQX were added to NMDG- and HEPES-ACSF.

### Quartz-seq2

#### Single-cell preparation

We used 6 mice (3 males and 3 females of Calb2-Cre/+; Ai3/+, C57Bl6/J background, age: 118-151 days old) for scRNAseq. After slicing brains, we followed the single-cell preparation protocol previously described (Hempel et al., 2007). Slices were incubated in bubbled HEPES-ACSF with 1 mg/ml pronase for 70 minutes at room temperature. PVT was isolated with a scalpel under a fluorescent dissection microscope. The isolated tissues were transferred to 1 % FBS/HEPES-ACSF and triturated with a series of fire-polished Pasteur pipettes with different tip diameters (600 µm, 300 µm, and 150 µm). To label nuclei and dead cells, 1 µg/ml of Hoechst 33342 (Thermo Fisher) and 7-AAD (Becton Dickinson) were added to the cell suspension. FACS-ARIA II (Beckman Coulter) fluorescent cell sorter was used to sort live single cells (Hoechst +, 7-AAD -) to 384-well plates loaded with 1 μL of lysis buffer (0.12 mM dNTP mix, 0.3% NP-40, 1 unit/μL RNasin plus) and 0.1111 μM respective primers for reverse transcription (v32_384p01, UMI sequence – cell barcode -oligo dT).

#### Whole transcript amplification

The cell plate was heated at 70°C for 90 seconds for cell lysis and then kept 35°C for 15 seconds for RT primer hybridization. RT premix (1 μl: 2× Thermopol buffer, 1.25 units/ μL SuperScript III, 0.1375 units/μL RNasin plus) was added to each well, and RT was performed at 35°C for 5 min and 50°C for 50 min and stopped at 70°C for 15 min. The plate was inversely placed on a disposable collector and centrifuged to collect solution in each well. The pooled solution was cleaned with DNA Clean and Concentrator-5 kit (Zymo Research) and extracted with 20 μL water. TdT solution (25 μL : 1× Thermopol buffer, 2.4 mM dATP, 0.0384 units/μL RNase H (Thermo Scientific), 26.88 units/μL terminal transferase (Roche)) was added and poly(A) tailing reaction was performed at 37 °C for 75 s. For primer tagging, PCR I premix (11 μL :1.08492× MightyAmp Buffer version 2, 0.06932 μM Tagging primer, 0.05415 units/μL MightyAmp DNA polymerase) was added, then incubated at 98°C for 130 sec, 40°C for 1 min, then heated at 0.2°C/sec, and then incubated for 1min at to 68°C. PCR II premix (50.232 μL: 0.99697× MightyAmp Buffer version.2, 1.8952 μM gM primer) was added to 56.16 μL PCR I solution, and PCR was performed by 11 cycles of 98 °C for 10 s, 65 °C for 15 s, and 68 °C 5 min. The solution was supplied with 88 μL of 3 M sodium acetate (pH 5.2) and 17.6 mL of PB-Buffer (Qiagen) and purified with MinElute Spin Column (Qiagen) to 32 μL of nuclease-free water. The solution was subsequently purified with 32 μL of Ampure XP beads (Beckman Coulter).

#### Library construction

the amplified cDNA (5 -10 ng) was added to 130 μL of nuclease-free water and placed in a tube with AFA Fiber. DNA fragmentation with LE220 (Covaris) was performed with a condition of 15 % duty factor, 450 W peak incident power, 200 cycle per burst for 80 second. The fragmented DNA was purified with DNA Clean and Concentrator and eluted to 10 μL of nuclease-free water. End-Repair mix (2 μL: 1.4 μL of End repair & A-tailing buffer and 0.6 μL of End repair & A tailing Enzyme (KAPA Biosystems)) was added to the fragmented DNA, and an end-repair reaction was performed by incubating at 37°C for 60 min and 65 °C for 30 min. Adaptor buffer (2 μL :1.5 μM truncated adaptor, 10 mM Tris-HCl pH 7.8, 0.1 mM EDTA pH 8.0, 50 mM NaCl) and ligation premix (8 2 μL : 6 μL of ligation buffer, 2 μL of DNA ligase (KAPA Biosyste))were added to the solution, and ligation reaction was performed at 20°C for 15 min. The solution was purified with 18 μL Ampure XP beads and extracted with 22 μL nuclease-free water. PCR premix (32 μL :25 μL of 2×KAPA HiFi ready mix, 1.75 μL of 10 μM TPC2 primer, 10 μM P5-gMac hybrid primer) to 18 μL of adaptor-ligated cDNA, and PCR was performed with 8 cycles of 98 °C for 15 s, 60 °C for 30 s, and 72 °C for 1 mim and following incubation at 72 °C for 5 min. Finally, the library was purified with 40 μL Ampure XP beads and eluted with 20-30 μL nuclease free water.

#### Sequencing

The indexed libraries were sequenced with NextSeq 500 (Illumina). Sequencing cycles were performed as following: Read1, 23 cycles; Index1, 6 cycles; Read2, 63 cycles.

#### Data Analysis

The Quartz-seq2 pipeline (https://github.com/rikenbit/Q2-pipeline) was used to generate a digital expression matrix from BCL files. For transcript mapping with RNA-star (Dobin et al., 2013), we used a custom index file made from a reference mouse genome (mm10/GRCm38) and transgenes (EGFP and Cre). For expression data analysis, we used Scanpy (https://scanpy.readthedocs.io/) python package and followed the standard analysis procedure described in the tutorial. The “raw” counting data were normalized to a million per cell. The Louvain algorithm in the Scanpy package and following Single-Cell Clustering Assessment Framework (Sccaf 0.0.10) were used for clustering and evaluation of clustering accuracy (Miao et al., 2020). The quasi-likelihood F-test function of edgeR (Chen et al., 2016) was used for differentially expressed gene (DEG) analysis between cell types. For gene network analysis, we followed a pySCENIC protocol (Van de Sande et al., 2020) and used Cytoscape (https://cytoscape.org) for network visualization.

### *in situ* hybridization

#### Multicolor in situ hybridization

RNAscope Multiplex Fluorescent Reagent Kit v2 (Advanced Cell Diagnostics) and Opal 520, 570, and 690 fluorophore reagent packs (AKOYA bioscience) were used for multicolor *in situ*. Unfixed, fresh brains from both sexes of C57Bl6/J mice were embedded in OCT compound (Sakura Finetech, Japan) and frozen sections were made at 16 μm. Sections were mounted on MAS-coat slide glasses (Matsunami, Japan), stored in a sealed box, and kept at -80°C until use. We followed the manufacturer’s protocol but skipped the protease treatment step since it was deleterious to the sections.

#### Imaging and signal processing

Images were taken with TCS SP8 confocal microscope (Leica) at 16-bit depth per channel. Fiji and custom Jython scripts were used for image analysis and quantification. Briefly,1) the PVT area was manually selected to make an ROI mask. 2) A DAPI image was cropped with the mask. 3) After background subtraction, the image was binarized. 4) ROIs around nuclei were set with the particle analyzer plugin then manually corrected. 5) The ROIs were overlayed to other channels, and mean signal intensity of each ROI were quantified. GaussianMixture function from Scikit-learn was used to estimate noise and signal curves. To clip signals for dot plots in Figure 2F and G, each signal value below the threshold of corresponding was set to the threshold.

### AAV preparation

pAAV-CAG-Flex-EGFP (#59331), pAAV-Ef1a-DO-hChR2(H134R)-mCherry-WPRE-Pa (#37082), and AAV-hSyn-DIO-hM3D(Gq)-mCherry (#44361) were obtained from addgene. For each AAV preparation, the 293T cells (obtained from RIKEN Bioresource Center, RCB2202) were seeded to 5 to 10 of Φ 15 cm dishes at 1.2-1.5 × 10^7^ cells/dish. An AAV transfer vector (6 µg/dish), pHelper (Agilent, 10 µg/dish), and AAVDJ/8 (Cellbio labs, 6 µg/dish) were mixed in 1 ml PBS/dish and supplied with 88 µl of 1 µg/ml polyethyleneimine (PEI MAX, Polyscience). After 5-10 minutes of incubation, the plasmids-PEI mixture was applied to the dishes. Culture media were harvested 3 days after transfection. After centrifugation at x 3,000 g for 20 minutes and filtration with a 0.25 µm filter, the media were loaded to thick-wall polypropylene tubes at 30 ml/tube. Being loaded sucrose cushion (2 ml 20 % sucrose/PBS) to the bottom, tubes were ultracentrifuged at 20,000 rpm with SW-28 (Beckman Coulter) rotor for 2 hours at 20 C. After the removal of supernatants, the pellets were dissolved with 1 mM MgCl_2_ / PBS. The AAV solution was dialyzed and concentrated with Amicon Ultra 15 centrifuge filter (100 kD, Millipore) and subsequently aliquoted and stored at -80°C.

### Stereotaxic injection

Mice were deeply anesthetized with a mixture of medetomidine (0.75 mg/kg), midazolam (4.0 mg/kg), and butorphanol (5.0 mg/kg) and then mounted on stereotaxic stage (SR-6M-HT and SMM-200, Narishige, Japan). After application of hair-removal cream and eye ointment, the skull was exposed, and heights of the bregma and the lambda were leveled. Grass capillaries (Wiretrol II, Drummond, USA) were pulled with a puller (P-97, Sutter Instrument, USA) to make needles. A needle was backfilled with mineral oil, and then attached to our custom injector made with MO-10 manipulator (Narishige, Japan). Virus solution was loaded from the tip of the needle. A hole (∼ Φ 1.5 mm) on the skull was made with a dental drill. To avoid breeding from the superior sagittal sinus, the needle tip was inserted slightly away from the sinus and moved laterally to the midline. The needle was inserted at a corresponding injection site (for aPVT: 0.4 mm posterior and 3.8 mm ventral from the bregma, for pPVT: 1.75 mm posterior and 3.05 mm from the bregma) and virus solution (100-200 nl) was injected at 50 nl/min. The needle was taken after waiting for 2 minutes. Xylocaine jerry was applied on the suture. For optical fiber implantation, the manipulator was set to 6 degrees from the vertical axis, and a cannula (in-house made, fiber: FT200EMT (Thorlab), ferrule: CFLC230 (Thorlab)) was inserted 200 μm above from the injection site (for aPVT: 0.4 mm posterior, 0.398 mm lateral, and 3.62 mm ventral, for pPVT: 1.75 mm posterior, 0.33 mm lateral, and 2.85 mm ventral from the bregma).

### Whole-brain imaging with TissueCyte

#### Surgery

Mice were deeply anesthetized with a mixture of medetomidine (0.75 mg/kg), midazolam (4.0 mg/kg), and butorphanol (5.0 mg/kg) and perfused with PBS and following 4 % paraformaldehyde in PBS. The brains were fixed in 4 % paraformaldehyde/PBS in 4°C for overnight and transferred to PBS.

#### Image acquisition

Block preparation (brain fixation, oxidization of agarose, brain embedding, and block crosslinking) was performed as described (Ragan et al., 2012). The block was washed with PB and kept dark at 4C until imaging. Tissuecyte 1400FC (TissueVision) equipped with Chameleon Ultra II laser (Coherent) and a 16x water-dipping objective (Nikon MRP07220, NA 0.8) was used for serial sectioning and imaging. The laser wavelength was tuned at 920 nm. Brains were coronally sectioned with a vibratome at 50 μm with a frequency of 60 Hz and a speed of 0.5 mm/sec, and each mosaic image (832 × 832 pixel, resolution at 1.382 μm) was taken at 50 um from the cut surface.

#### Image stitching and intensity correction

Vignetting artifacts that appeared as decreasing intensity from tile centers to their boundaries were reduced using our in-house intensity correction software (Skibbe et al., 2019). Tiles were stitched to entire 2D images of the coronal sections. The positions of tiles were determined by the 3D world coordinates that were provided by the TissueCyte microscope. Since the image quality was low or partially corrupted at the boundaries of an image tile, 20 pixels from the left and right side, and 30 pixels from the top and the bottom of each tile were cropped. That still gave an about 20 pixels margin for overlapping tiles during stitching. Overlapping parts of the image tiles were merged using linear blending.

#### Full resolution and low-resolution image stacks

The entire brain image stacks were reconstructed in full resolution. With a spatial resolution of 1.34 × 1.34 × 50 μm^3^ a stack consisted of about 321 coronal sections, each containing about 8,886 × 6,912 pixels. Coronal sections were stored as 16-bit png image files (lossless). The full resolution image stacks were scaled down to 25 × 25 × 50 μm^3^. Those image stacks were used for image registration and GFP signal segmentation. The images were stored in the NIFTI file format.

#### Image registration

All 3D image stacks were mapped to the Allen Mouse Brain Common Coordinate Framework version 3 (Wang et al., 2020) that was downloaded from the Scalable Brain Atlas website (https://scalablebrainatlas.incf.org/mouse/ABA_v3#downloads). The data contained a population average mouse brain image template based on the auto-fluorescent background signal of TissueCyte image stacks. It further comprised brain region annotations. For the mapping, the ANTs registration toolkit was used (Supplementary Figure4G) (Bakker et al., 2015). A first step determined the mapping between a 3D image stack of channel 1 of an individual brain and the Allen Mouse Brain template image. Once the transformation was computed, it was used to map the image stacks of the remaining channels. After mapping, all image stacks shared the same image space.

#### Segmentation and normalization of signal

The TissueCyte system we used has 4 detectors for multicolor imaging, and we utilized “bleed-through” signals for background subtraction and estimation of injection sites. Briefly, the background signal was reduced by subtracting the darkest channel from the brightest channel. Next, the signals were normalized between the interval [0,1]. Since the signal directly in the injection was saturated, we did not use it as maximum for normalization. Instead, we used a heuristic to find the brightest intensity value in its proximity. The volume of the injection site was determined by thresholding channel 1 at a value that was half of its maximum intensity. A connected component analysis identified the largest volume in the result. This step was necessary to remove falsely segmented bright regions other than the injections site. Morphological dilation was applied to slightly increase the volume of the injection site. In that way, we identified the volume close to the injection site from which we picked the maximum intensity for normalizing the reporter signal was drawn.

### Electrophysiology

We use adult 90-180 days old adult mice of both sexes. A brain slice was submerged in ACSF (124 mM NaCl, 2.5 mM KCl, 1.2mM NaH_2_PO_4_, 24 mM sodium carbonate, 5 mM HEPES, 12.5 mM Glucose, 310 mOsm, pH 7.3-4) at 32 °C. Synaptic blockers (50 µM D-AP5, 20 µM CNQX, and 20 µM picrotoxin) were added to ACSF for recording intrinsic properties. Recording pipets (resistance 2.2 – 4 MΩ) were filled with the internal solution (20 mM KCl, 100 mM Kgluconate, 10 mM HEPES, 4 mM Mg-ATP, 0.3 mM Na-GTP, 10 mM Na-phosphocreatine). Since PVT neurons’ activities have diurnal variance (Kolaj et al., 2012), we performed recording at a fixed schedule; brains were dissected around 10 AM, and recordings started from around 12 PM. A set of recordings were performed with ACSF, and second set of recordings were performed with ACSF containing one of ligand. Since ligand-induced depolarization of resting membrane potential could last more than 30 minutes after changing solution back to regular ACSF, new brain slices were used in every recording session. Signals were collected with Multiclamp 700B and Digidata 1440 (Molecular Devices). pClamp 10 (Molecular Devices) was used for data collection. Data were analyzed with jupyter notebook (https://jupyter.org) and pyabf (https://swharden.com/pyabf/).

### Food intake test

We followed a protocol described in (Zhang and van den Pol, 2017). Each mouse (equal numbers of males and females) was isolated 3 days before the test and had ad libitum access to water and the mouse chow (CRF-1, Charles River). All tests were performed between 1 PM to 3 PM. CNO (1mg/kg) or saline was injected 30 minutes before the session. After the session, the mice were returned to their home cages and chows were weighed. Second tests were performed in the next week with the same method, but the other solution was injected.

### Real time place preference test

An optical fiber attached with a blue DPSS laser (473 μm, 150 mW, Shanghi Laser & Optics Centure Co., China) was connected to an implanted cannula. The laser power was adjusted to 8 -10 mW at the tip of a cannula. A mouse was placed in a chamber with two connected rooms (30 × 60 cm, 10 cm opening). After 20 minutes of adaptation, ten minutes each of recording sessions started without laser stimulation (first session: baseline recording). In the second session, the laser stimulation (20 Hz) was triggered when the mouse was in one of the rooms, and the laser-on room was switched to the other room in the third session. RaspberryPi 3 and a custom python program were used in camera recording and laser stimulation control.

## Data and code availability

scRNAseq data (fastq files and digital expression matrix) are publicly available in the DDBJ Sequenced Read Archive under the accession numbers DRA013342 and E-GEAD-470. Original codes in this study has been deposited at GitHub (https://github.com/yshima/2022_PVT_celltype_paper).

